# Interval between two sequential arrays determines their storage state in visual working memory

**DOI:** 10.1101/2020.01.08.814889

**Authors:** Ziyuan Li, Jiafeng Zhang, Tengfei Liang, Chaoxiong Ye, Qiang Liu

**Author notes:** These authors contributed equally to this work and should be considered co-first authors (Ziyuan Li, Jiafeng Zhang). Corresponding author: Qiang Liu, Ph.D., Professor, Institute of Brain and Psychological Sciences, Sichuan Normal University, Chengdu, China, 610000.

## Abstract

The visual information can be stored as either “active” representations in the active state or “activity-silent” representations in the passive state during the retention period in visual working memory (VWM). Catering to the dynamic nature of visual world, we explored how the temporally dynamic visual input was stored in VWM. In the current study, the memory arrays were presented sequentially, and the contralateral delay activity (CDA), an electrophysiological measure, was used to identify whether the memory representations were transferred into the passive state. Participants were instructed to encode two sequential arrays and retrieve them respectively, with two conditions of interval across the two arrays: 400ms and 800ms. These results provided strong evidence for the state-separated storage of two sequential arrays in different neural states if the interval between them was long enough, and the concurrent storage of them in the active state if the interval was relatively short. This conclusion was valid only when the participants encountered the task for the first time. Once participants have formed their mindset, they would apply the same storage mode to the subsequently extended or shortened interval condition.

## Introduction

The visual working memory (VWM) system is responsible for the transient retention and manipulation of visual information from the dynamic visual environment^1,2,3^. It helps to bridge across changes and create temporal continuity from multiple, successively occurring displays in the service of advanced cognition. A fundamental property of this system is its limited capacity. Recent studies consider that approximately four objects can be retained at once in VWM, and this information is encoded in the form of integrated visual features, rather than as a collection of disconnected visual features^4,5,6^. These studies usually employ static displays to present all stimuli simultaneously. However, visual events evolve over time in many cognitive activities. Concerning the dynamic nature of the visual environment, investigating VWM via sequential arrays could comprehensively explain the representations from the real world.

Recently research efforts have focused on dynamic visual processing to make great advances in scientific knowledge about memory. For example, Jiang and Kumar explored whether the two sequentially presented arrays, separated by a variable interstimulus interval (ISI), were represented as an integrated image or two separated images in visual short-term memory (VSTM). They proposed that two sequential visual arrays were held separately at an ISI of 500 ms or longer^7^. Subsequently they employed the same paradigm to further explore the dynamics of visual information, suggesting that VSTM had a constant capacity independent of the stimulus onset asynchrony (SOA)^8^. Additionally, Gorgoraptis el at. designed sequential and simultaneous presentation condition to examine the dynamics and flexibility of WM resources. Their study revealed that WM resources could be dynamically and flexibly updated as new items had to be stored, but redistribution of resources with the addition of new items was associated with misbinding object features^9^. Notably, these studies adopted a combined probe task in which mnemonic representations would be all or randomly probed in a single probe array. And the specific task potentially compelled participants to combine all memory information in their mind. This processing manner corresponded well to the dynamic cognitive activities, such as chess game which requires updating of information gathered from many instances and information-retention across space and time. Nevertheless, dynamic visual processing also involves non-combined retrieval. For example, a learner models a dance following a dancer’s performance. Because the dance is a particular series of graceful movements, the learner must remember these dynamically appeared actions, and then performs them serially and accurately. Thus, the dynamics of visual information should endow not only the presentation of memory stimuli, but also the retrieval of mnemonic representations. Here, we make efforts to explore how the two sequential arrays are stored in VWM using a sequential-encoding-sequential-retrieval task in which two memory arrays are encoded sequentially and then retrieved sequentially in the same order.

The multi-state storage mechanism of VWM has been extensively recognized. One form of evidence comes from “activity-silent” synaptic mechanisms by which humans can hold information in WM^10^. The “activity-silent” model proposed that the memory representations could be stored in either “active state” that was accompanied by persistent delay activity, or “passive (or silent) state” that could not be directly detected by recording techniques, but still maintained the relevant information^11^. In a memory-guided saccade task (MGS), it could be observed a “ramp-up” that indicated the robust activity emerging only at the late delay period for upcoming response cue, as well as a silent moment during which the robust delay activity was absent between the memory stimuli onset and WM-behavioral response^11,12^. Similarly, when adding a attention task demand to the classic MGS task, WM-specific delay activity was abolished during the dual task period. But once the attention task was completed, the WM delay activity was “reawakened”, presumably in time for WM-guided behavior^11,13^. The two studies exactly supported the “activity-silent” model, showing that persistent neural activity was not critical for the maintenance of WM content but instead reflected task relevance of memoranda. Thereby, it could be known that whether the memory representations were maintained in the active state or the passive state was determined by the current task relevance. According to the “activity-silent” model, it could be reasoned that, in the sequential-encoding-sequential-retrieval task, the representations of currently task-irrelevant array would be held in the passive state, while the others should be held in the active state for the current encoding or the imminent probe.

Within the field of electroencephalography, researchers find a sustained contralateral negativity over posterior electrode sites during the retention period of VWM tasks through the event-related potential (ERP) studies. It has been proven that the amplitude of contralateral delay activity (CDA) increases as the number of memory items increases, up to the individual’s working memory capacity limit^14,15^. CDA is widely used as an effective marker to identify the number of items in the active state^16^. That is, the CDA amplitude would decrease quickly and disappear when the memory representations are transferred from the active state to the passive state. In addition, the CDA is not modulated by perceptual requirements and the number of currently attended locations^17,18,19,22^. Hence, if the CDA is applied to the dynamic sequential-encoding-sequential-retrieval task, then we can explicitly reckon whether the first array is transferred into the passive state during the maintenance period in VWM.

The sequential-encoding-sequential-retrieval task is rarely exploited to explore the storage mode of sequential arrays, but it has been adopted to determine what aspect of WM performance the CDA reflects^17^. However, Ikkai et al.’s results suggested that the CDA amplitude during the encoding of the second array doubled compared to that following the first array, which implied that the two sequential arrays were concurrently stored in the active state following the second array. The results observed by Ikkai et al. do not suggest that WM representations of the first array are transferred into the passive state (i.e. activity-silent maintenance).

The brevity of the interval between two sequential arrays might be a plausible account of this contradiction. The interval between two arrays is a critical factor to determine whether the two sequential arrays are separately retained. Jiang and Kumar conducted a paradigm in which two dot arrays, separated by an ISI of 0, 200, 500, or 1,500ms, were sequentially presented. Though they mainly investigated the representation mode (an integrated representation or two separated representations) of the two arrays regardless of their storage state, and the memory task for spatial configuration of multiple dot locations was completely different from remembering the color of squares in Ikkai et al.’s study, their findings at least showed that the relatively long interval between two arrays could make the leading array consolidated well and further determine the separation of the two sequential arrays. However, in Ikkai et al.’s study, the interval between two memory arrays was only 400ms. So the items of the first array were not fully consolidated, and then they were continuously retained in the active state, together with the second memory array. Therefore, it could be deduced that extending the interval between the two sequential arrays would achieve the goal that the currently task-irrelevant representations were transferred into the passive state from activity in VWM, resulting in a state-separation of the two dynamic arrays.

In the current study, we adopted the sequential-encoding-sequential-retrieval task to examine whether the storage mode of two sequential arrays was modulated by the interval retention between them in VWM, and the CDA was employed to identify the storage state of VWM representations. According to the reasoning above, we predicted that if the interval between the two memory arrays was extended enough, the two arrays would be separately stored in active and passive states; In contrast, if the interval still stayed short as Ikkai et al.’s experiment 2, then we would replicate the results that the two arrays were concurrently stored in the active state during the encoding of the second array.

## Method

There were two experimental conditions as depicted in Figure 1: short interval condition (400ms) and long interval condition (800ms). We aimed to test whether participants could potentially succeed in the two interval conditions by employing concurrent maintenance of the two arrays in the active state, or state-separated maintenance in different neural states. If the CDA amplitude following the second array did not increase compared to that following the first array in the long condition (800ms), we could confirm that the two arrays were stored in different neural states due to the state transformation of the first array; If the CDA amplitude did increase following the second array compared to the first array when confronting the relatively short interval period (400ms), we then deduced that the two arrays were concurrently stored in the active state during the encoding of the second array. The current experiment was a within-subjects design. Participants were instructed to complete two conditions (short and long) with 4 blocks each. The order of condition was counterbalanced across subjects. They needed to perform 8 blocks consisting of 512 trials in total.

**Figure 1:**
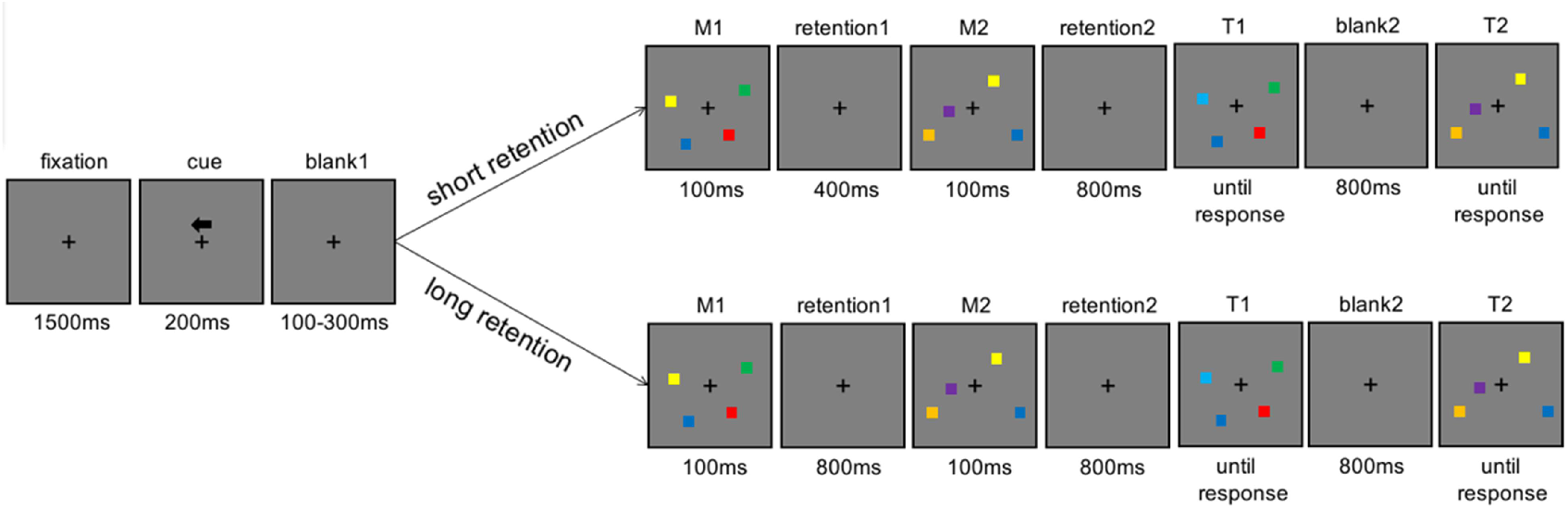
Trial schematics for the short condition (upper) and the long condition (lower).

### Participants

Twenty-two participants (8 males) from Liaoning Normal University voluntarily took part in the experiment. The averaged age was 21 ranging from 18 to 26. All participants were neurologically normal and reported a normal or corrected-to-normal vision without color blindness. They participated in the experiment with a compensation of CNY 50. They provided written informed consent when they arrived at the lab. The experiment would last about 2.5 hours. The experiment was carried out in accordance with the Declaration of Helsinki and approved by the institutional ethics committee of the Liaoning Normal University. Two additional participants recruited in the experiment were excluded from the further analysis: one participant because of the excessive eye-blink or eye-movement artifacts (in excess of 25%) and one participant because of the poor performance (lower than the chance level). No participants were excluded due to high trial rejection rates (< 25% of trials remained). The averaged rejection rate per subject was 23%, and SD was 11% (an average of 197 trials for each condition were analyzed).

All participants were divided into two groups according to the order of experimental condition: those (11 participants) who performed the short condition first and then the long condition were called short-long group, and those (11 participants) who performed in converse order were called long-short group.

### Stimuli and procedure

In the experiment, the stimuli were presented with E-prime 2.0 on a CRT screen in a brightly-lit room. The schematic of a trial is illustrated in Figure 1. All stimuli were presented on the gray background. Participants were instructed to fixate the fixation cross from 70 cm of viewing distance. The fixation cross was presented in the middle of the screen throughout the whole trial. A trial began with a 1500ms-retention, and then a arrow cue was presented above the fixation cross for 200ms, instructing the participants to remember the colors of squares in the cued field from the two arrays. After a random retention (100ms-300ms), two memory arrays (M1 M2) appeared sequentially with a 400ms-interval in the short condition and an 800ms-interval in the long condition. After an 800ms-interval retention following the M2, the first test array (T1) and the second test array (T2) were also presented sequentially. At the onset of the test arrays, participants responded whether the test arrays (T1 T2) were identical to the memory arrays (M1 M2) respectively. If they were identical, participants were instructed to press the “F” in the keyboard, or press ‘J’. The test arrays did not disappear until the participants made response. Through all trials, participants were informed to make a button press as accurately as possible, without emphasis on the speed. Memory and probe items were presented within 4 × 7.3° invisible regions bilaterally, centered 3° to the left and right of the middle of the screen. Colored squares (0.65 × 0.65°) were randomly chosen from a set of seven discriminable colors (red, orange, blue, yellow, green, indigo and purple) without replacement within a trial. Item positions were randomized in these trials with a constraint that no square was presented within 2° with each other^17^. Participants had a rest of at least 30s between blocks. They would be informed of what the interval condition was prior to each condition. Before the formal experiment, all participants were asked to practice at least 16 trials per condition and reached 75% accuracy to understand the experimental procedure well.

### Electrophysiology (EEG) recording and analyses

The EEG signals were recorded by using a 64-channel amplifier. ANT Neuro EEGO mounted in a cap using 10/20 montage, including Fp1, Fp2, Fpz, AF3, AF4, GND, AF7, AF8, F1, F2, F3, F4, F5, F6, Fz, FT7, FT8, FC1, FC2, FC3, FC4, FC5, FC6, FCz, T7, T8, C1, C2, C3, C4, C5, C6, Cz, TP7, TP8, CP1, CP2, CP3, CP4, CP5, CP6, CPz, P1, P2, P3, P4, P5, P6, P7, P8, Pz, PO3, PO4, PO7, PO8, POz, O1, O2 and left and right mastoid electrodes. In these electrodes, CPz served as the on-line reference and GND served as the ground electrode. The off-line signals were analyzed with EEGLAB v13.5.4b in MATLAB R2016a, and the data were only filtered with a low-pass of 40 Hz. The average of the left and right mastoids was the re-reference. The electrodes were placed 1 cm to the left and right of the external canthi to record horizontal eye movement (HEOG). All the electrode impedances were kept below 10 kV and the data were collected at a sampling rate of 500 Hz.

The continuous signals were segmented from 100ms before to 1400ms (in the short condition) or 1800ms (in the long condition) after the first memory array onset. The EEG data were baseline-corrected with the 100ms prior to the first memory array onset. The EOG artifacts were corrected by independent component analysis algorithm^23^. Those trials containing the artifacts of blinks and eye movements (exceeding ±80μV) were excluded from further analysis. Participants with excessive EEG artifacts would be excluded from the analyses. Bad channel was replaced by the interpolation method. The CDA was calculated with the mean amplitude of difference wave (contralateral - ipsilateral), averaged over electrode sites P7 and P8^24^. The CDA mean amplitude was analyzed with two time windows between 300ms and 500ms after the first memory array onset and the second memory array onset ^17^. Bonferroni correction was used for multiple comparisons.

## Results

### Behavioral results

Statistical analyses were computed using SPSS. 21. The Shapiro-Wilk test was adopted to testify the normality of the data in each condition (short condition, test 1: p=0.233; short condition, test 2: p=0.119; long condition, test 1: p=0.137; long condition, test 2: p=0.569). A 2 (short condition vs. long condition) × 2 (test 1 vs. test 2) repeated measures ANOVA was conducted to analyze the behavioral accuracy. The main effect of condition was significant (F (1, 21) =16.088, padj=0.001, *η_p_*^2^ =0.434). The main effect of test was also significant (F (1, 21) =55.450, padj< 0.001, *η_p_*^2^ =0.725). There was no significant interaction between condition and test (F (1, 21) =1.402, p=0.250, *η_p_*^2^ =0.063). In the short condition, the mean accuracy of test 1 and test 2 were 88.1% and 75.2%, and SD were 0,059 and 0.079, respectively. In the long condition, the mean accuracy of test 1 and test 2 were 90.7% and 79.6%, and SD were 0.047 and 0.077, respectively. The long interval between two arrays could promote the behavioral performance. Additionally, the accuracy of test 2 was lower than that of test 1 irrespective of the interval, which might be accounted from the interference of making response prior to the test 2, as well as the decay over time.

Considering that two distinct groups have been divided according to the order of experimental condition, we examined the behavioral performance separately for the short-long group (completed the short conditions before the long conditions, N=11) and the long-short group (completed the short conditions after the long conditions, N=11). Firstly, we conducted a mixed-design ANOVA of condition (long condition vs. short condition) and test (test 1 vs. test 2) as within-subject factors, and group (short-long group vs. long-short group) as a between-subject factor, suggesting that there were no significant interactions between condition × test × group (F(1, 20)=0.395, p=0.537), condition × group (F(1, 20)=0.006, p=0.940), and test × group (F(1, 20)=1.475, p=0.239). The results indicated that the short-long group and the long-short group did not differ in their behavioral performance. For the short-long group, a 2 (short condition vs. long condition) × 2 (test 1 vs. test 2) repeated measures ANOVA showed that the main effect of condition was significant (F (1, 10) =8.361, padj=0.016, *η_p_*^2^ =0.455). The main effect of test was also significant (F (1, 10) =27.292, padj<0.001, *η_p_*^2^ =0.732). There was no significant interaction between condition and test (F (1, 10) =0.106, p=0.751, *η_p_*^2^ = 0.011). In the short condition, the mean accuracy of test 1 and test 2 were 87.8% and 77.3%, and SD were 0.071 and 0.072, respectively. In the long condition, the mean accuracy of test 1 and test 2 were 90.8% and 81.2%, and SD were 0.050 and 0.064, respectively. While for the long-short group, a 2 (short condition vs. long condition) × 2 (test 1 vs. test2) repeated measures ANOVA showed that the main effect of condition was significant (F(1,10)=7.119, padj=0.024, *η_p_*^2^ =0.416). The main effect of test was also significant (F(1, 10) =30.125, padj<0.001, *η_p_*^2^ =0.751). There was no significant interaction between condition and test (F (1, 10) =2.524, p=0.143, *η_p_*^2^ =0.202). In the short condition, the mean accuracy of test 1 and test 2 were 88.5% and 73.2%, and SD were 0,048 and 0.084, respectively. In the long condition, the mean accuracy of test 1 and test 2 were 90.6% and 78.0%, and SD were 0.045 and 0.089, respectively. Figure 2 shows the behavioral accuracy across all participants (top), the short-long group (bottom left) and the long-short group (bottom right) in each condition.

**Figure 2:**
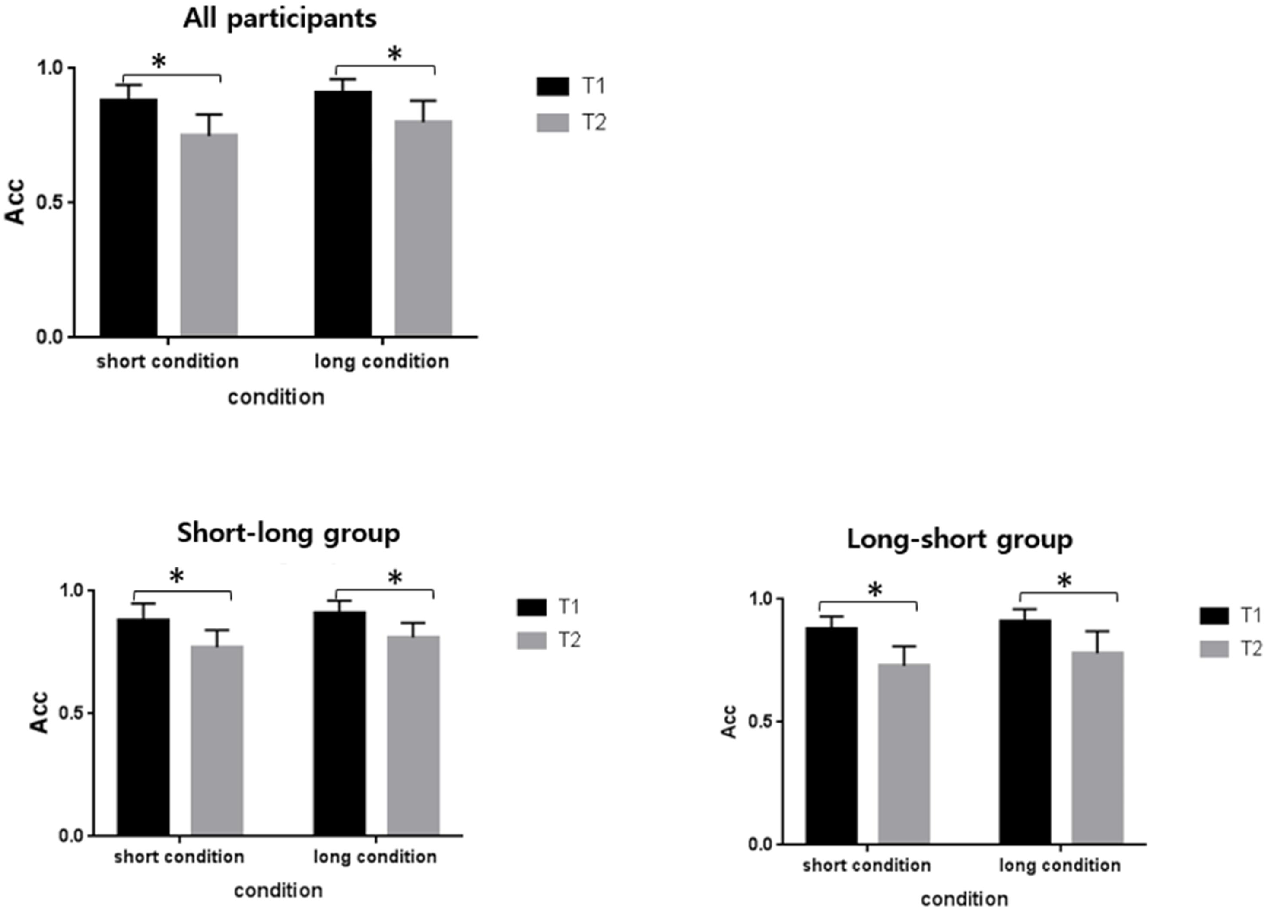
Behavioral accuracy of the test 1 and test 2 across all participants, the short-long group and the long-short group in each condition. The error bars indicate the standard deviation.

### Electrophysiological results

CDA amplitudes in the short and long conditions are plotted across all participants in Figure 3A. Strong negative wave arose over posterior electrode sites about 250ms after the memory arrays onset during the retention period. By visual inspection, CDA amplitude appeared much larger following the second memory array than the first memory array in the two conditions. The Shapiro-Wilk test was adopted to testify that the data in all single conditions were normally distributed (short condition, early delay: p=0.061; short condition, late delay: p=0.349; long condition, early delay: p=0.252; long condition, late delay: p=0.080). The analysis of CDA amplitude was conducted by a 2 (short condition vs. long condition) × 2 (early delay vs. late delay) repeated measures ANOVA (early delay: 300–500ms following the first memory array onset; late delay: 300–500ms following the second memory array onset). Figure 4A shows the comparison of CDA amplitude during the two time windows in each condition. There was no main effect of condition (F (1,21)=1.157, padj=0.294, *η_p_*^2^=0.052), but the main effect of delay was significant (F(1,21)=8.767, padj=0.007, *η_p_*^2^ =0.295). The interaction of condition and delay was not significant (F(1,21)=3.322, p=0.083, *η_p_*^2^=0.173). The CDA amplitude increased during the late delay compared to the early delay independent of the interval retention.

**Figure 3:**
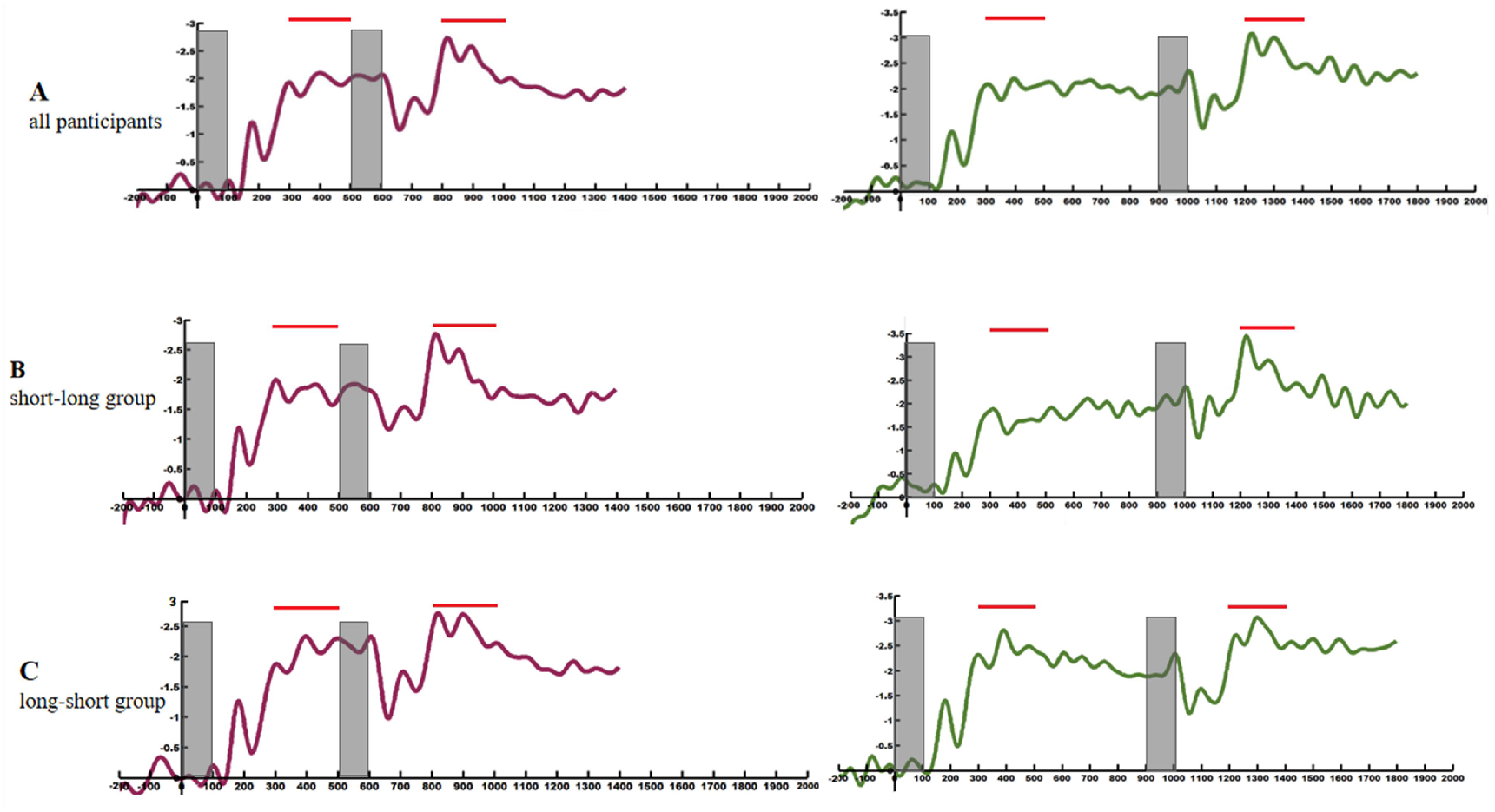
Event-related potential (ERP) data from the short condition (violet line) and the long condition (green line) averaged across all participants (top), the short-long group (middle), and the long-short group (bottom). Time-locked to the onset of the first memory array. Negative is plotted upward. The sites recorded in posterior lateral are P7\P8 to calculate the difference wave (contralateral-ipsilateral). The gray field indicates the presence of memory arrays. The red line on the top highlights the time window.

**Figure 4:**
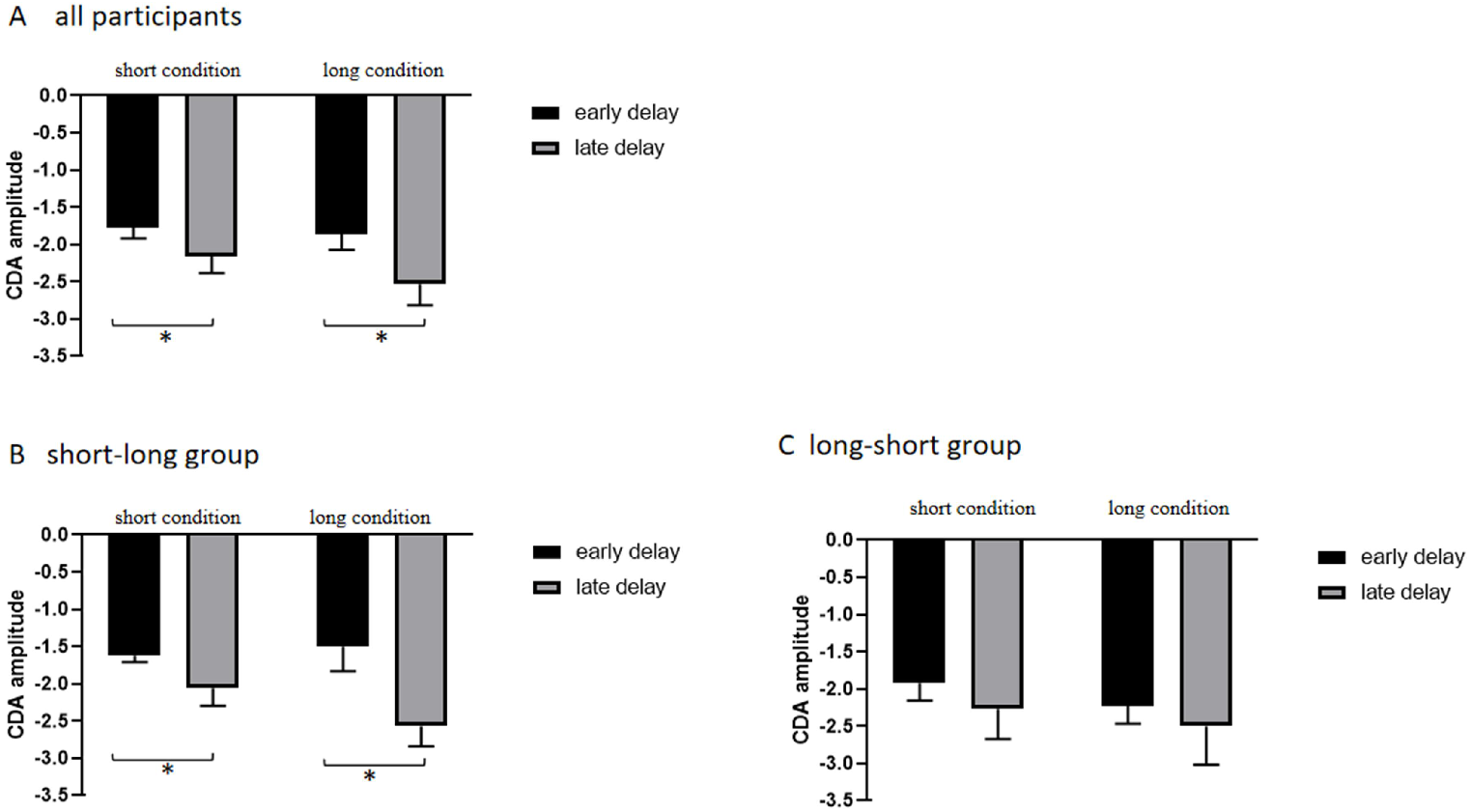
CDA amplitudes during the early delay (between 300ms and 500ms following the first memory array onset) and the late delay (between 300ms and 500ms following the second memory array onset) across all participants, the short-long group and the long-short group. The error bars indicate the standard error.

Notably, priming could greatly affect the interpretation of incoming input in VWM. The results of Balaban’s experiment 4 suggested that the manner of performing task that was primed before the main experimental phase could influence the interpretation that first appeared during the main experimental phase^21^. So it was necessary to take priming effect into consideration. Given that there were two conditions with a within-subjects design, and the order of condition was counterbalanced across subjects, it could be predicted that the CDA amplitude would have distinctly different trends in the short and long conditions when removing the order effect.

Firstly, the data of all participants were subjected to a mixed-design ANOVA of condition (long condition vs. short condition), delay (early delay vs. late delay) and group (short-long group vs. long-short group). The results revealed a significant interaction between these three factors (F(1, 20)=7.548, p=0.012, *η_p_*^2^ =0.274), suggesting that the short-long group and the long-short group were different from each other in EEG results. Across the short-long group, by visual inspection, the CDA amplitudes after the second array appeared much larger than that after the first array in the two conditions (Figure 3B). The CDA amplitudes during the two time windows were also analyzed by a 2 (short condition vs. long condition) × 2 (early delay vs. late delay) repeated measures ANOVA. The results showed that the main effect of condition was not significant (F(1, 10)=0.414, padj=0.535, ηp ^²^=0.040). But a significant main effect of delay was observed (F(1, 10) =36.672, padj < 0.001, ηp ^²^ = 0.786). The interaction between condition and delay was significant (F (1, 10) =24.338, p = 0.001, ηp ^²^ =0.709) (Figure 4B). Follow-up t-test run separately for trials in the short condition and the long condition. The Shapiro-Wilk test was used to verify the normality of the data (short condition: p=0.217; long condition: p=0.611). In the short condition, the mean amplitudes in the early delay and the late delay were −1.62 and −2.06, and SD were 0.632 and 0.688, respectively (t(10)=2.720, p=0.022, 95% CI=[0.079 0.799]). While in the long condition, the mean amplitudes in the early delay and the late delay were −1.50 and −2.57, and SD were 1.092 and 0.904, respectively (t(10)=9.297, p<0.001, 95% CI=[0.815 1.329]). These results demonstrated that the two arrays were concurrently stored like a perfect additivity in the active state after the second memory array, and at the same time, they were trending toward an integration as the participants completed more trials. In contrast, across the long-short group, CDA amplitude did not obviously increase during the late delay compared to the early delay (Figure 3C). The CDA amplitudes of the two time windows were also analyzed by a 2 (short condition vs. long condition) × 2 (early delay vs. late delay) repeated measures ANOVA. The results showed that the main effect of condition (F (1, 10) =0.700, padj=0.422, ηp ^²^ =0.065) and delay (F (1, 10) =0.871, padj=0.373, ηp ² =0.080) were not significant, nor was there a significant interaction between condition and delay (F(1, 10) =0.143, p =0.713, ηp ^²^ =0.014) (Figure 4C). In the long condition, the mean amplitudes in the early delay and the late delay were −2.228 and −2.496, and SD were 0.785 and 1.711, respectively. While in the short condition, the mean amplitudes in the early delay and the late delay were −1.913 and −2.267, and SD were 0.788 and 1.325, respectively. For the long-short group, the first array was retained in the passive state when the second array was being encoded in the active state irrespective of the interval period. By analyzing the two distinct groups, we found that the storage mode adopted in the latter condition was always the same as that in the initial condition for each group. Therefore, it could be known that priming indeed existed.

The current study attempted to test whether the two sequentially presented arrays were stored as two state-separated images in the long condition. To abolish the priming effect, we only explored the short condition across the short-long group and the long condition across the long-short group. The results from the short condition replicated the findings by Ikkai et al.: the CDA amplitude following the second array was significantly larger than that following the first array, suggesting that the two arrays were concurrently maintained in the active state. Whereas in the long condition, there was no significant difference in CDA amplitude between the two delays, which indicated that mnemonic representations of the first array were transferred into the passive state during the encoding of the second array, and thus the two memory arrays were stored as two state-separated images in different states. In other words, when removing the priming, the results of the current study were consistent with our hypothesis that the two sequential arrays would be stored separately when the interval between two sequential arrays was long enough.

In addition, considering the different results when processing the EEG data in a distinct way, the results indeed showed a priming effect of storage mode. For the long-short group, the CDA amplitudes following the first and the second memory arrays were not significantly different in neither the long condition nor the short condition; while for the short-long group, the CDA amplitude in the late delay was significantly higher than that in the early delay in both long condition and short condition. Specifically, the storage mode that was indeed primed before another interval condition trials began could influence the subsequent storage mode, no matter what the next interval condition was short or long. Participants of the short-long group still employed a concurrent manner to retain two sequential arrays in the active state in the long condition; similarly, participants of the long-short group even applied a separated manner to the short condition. To sum up, whether the storage mode of two sequential arrays was concurrent in the active state or state-separated also depended on their mindsets which were formed in the initial condition and played a dominate role in subsequent condition regardless of the interval was long or short

## Discussion

The primary goal of this study was to explore how VWM stored the memory information over a more dynamic, sequential input and retrieval. To accomplish this, the sequential-encoding-sequential-retrieval task was adopted, in which we modulated the interval (400ms vs. 800ms) between two memory arrays. And the CDA was employed to identify whether the WM representations were maintained in the active state or the passive state. The results showed that the CDA amplitude during the encoding of the second array was significantly larger than that following the first array in both short and long interval conditions across all participants, indicating that the two arrays were concurrently stored in the active state independent of the interval. Given that the current study was within-subjects design, and the order of experimental condition was counterbalanced across subjects, all participants were divided into two groups: the short-long group and the long-short group. For the short-long group, it was found that the CDA amplitude following the second array was significantly larger than that following the first array in both short and long conditions. This result suggested that the two arrays were concurrently maintained in the active state after the second array onset regardless of the interval condition. Whereas, for the long-short group, there was no significant difference in CDA amplitude between the early and late retention periods in the two conditions, which suggested that the two arrays were stored separately in different neural states regardless of the interval as well. Importantly, the results from the short-long group in the short condition and the long-short group in the long condition revealed that the two sequential arrays were concurrently stored in the active state when the interval was relatively short, but the two were separately maintained in different neural states when the interval was extended enough. That was valid only when the participants initially encountered the task. The behavioral results suggested that the two storage modes generated comparable VWM performance.

Therefore, this study provides strong evidence that the temporal context between two sequential arrays makes a great contribution to the storage mode. The state-separated maintenance of two sequential arrays in VWM may occur if the interval (e.g. not less than 800ms) across two memory arrays is sufficient to transfer the representations of the first array into the passive state. Whereas if the shortage of interval (e.g. no more than 400ms) prevents the storage transformation, the two temporally segregated arrays are concurrently maintained in the active state as one perfect additivity which still contains the independent component of each array.

Moreover, the current results have revealed the role of mindset in the storage mode through the data of the short-long group and the long-short group in the two conditions, manifested by the priming effect. The current results allow us to conclude that the storage mode at the early stage of the experiment would be fixed and stay the same even in subsequently different condition. That provides a plausible account of why the results of all participants are not consistent with our hypothesis. What’s more, it can be known that it is a top-down voluntary process rather than a stimulus-driven process for participants to decide whether to transfer VWM representations into the passive state. When the storage transformation could optimize cognitive resource allocation, participants would change the storage state of VWM representations. However, if the shortage of time restricts the storage transformation, then the favored strategy is keeping the VWM representations in the active state until making response. This kind of transformation strategy might become a mindset in similar cognitive processing. Once the participants have formed their mindsets which do not need to transfer the VWM representations to the passive state, the transformation do not occur anymore. Therefore, the mindset of storage manner also affects whether the two sequential arrays are concurrently stored in the active state.

What’s more, based on the characteristic of the CDA that reflects the number of memory items in the active state, it has been found that, in a sequential-encoding-simultaneous-retrieval paradigm, the two memory arrays would be concurrently stored if the probe array was spatially congruent with the memory arrays; whereas the first memory array would be retained in the passive state during the encoding of the second array if the items of memory arrays were spatially translated and interleaved in the probe array^16^. Thus, it has been deemed that the spatial compatibility of displays is also an important factor to determine whether the two sequential arrays are viewed as a single extended event or two independent episodes.

In the current study, we interpret the lack of CDA amplitude increase following the second array as that only the second array is retained in the active state due to the storage transformation of the first array. But there may be an alternative interpretation that the two arrays are integrated to form “one object”. If so, it is a must that, in the short interval condition, they are also integrated as “one subject” to be maintained after the second memory array onset, and thus the increase of CDA amplitude should be absent. However, the fact that the CDA amplitude increase was observed following the second array in the short condition denies the assumption of “one object”. Overall, the current results support the interpretation that, because the first array is transferred into the passive state, only the second array is maintained in the active state during the encoding of the second array in the long interval condition.

With an 800 ms delay between arrays, the current results have shown that the CDA never went away. The presence of the “active” CDA indicates the first array being constantly retained in the active state. Curiously, this finding raises a question why the items of the first array are not transferred into the passive state during the 800ms delay. Recent studies have verified that activity during memory delays might serve an important role in prioritizing mnemonic content in preparation for efficient WM-guided behaviour^11^. And the neural activity would disappear after making response. Thus it is reasonable that the mnemonic representations would be retained in the active state as much as possible to enhance the representations themselves before they are probed within a limited delay retention. Hence, the first array was constantly retained in the active state, and transferred into the passive state during the encoding of the second array when encountering the 800ms interval between the two arrays. Moreover, the 800ms-delay might be just sufficient to consolidate the first array fully and no more spare time to transfer it to the passive state. And thus the state transformation of the first array only occurs during the encoding of the second array, which do not possibly produce any cost of performance. However, further study is needed to explore whether the items of the first array are transferred into the passive state before the second array onset when an even longer interval is used.

VWM plays an important role in linking the preceding and following scenes. Beyond much understanding of VWM from the static displays in which all stimuli are represented simultaneously, we attempt to explore the storage mode of VWM representations by presenting robust temporal dynamics over a very rapid timescale. Individuals need constant updating of information gathered from multiple successive occasions, so it could be energetically expensive to hold memory information in the active state all the time, especially if WM is in near-constant use constructing and maintaining an up-to-date world model of the content of the environment^11^. Therefore, the neural economy may promote the state-separation of the two sequential arrays in VWM when encountering a relatively long interval. To sum up, whether the two sequentially presented arrays are concurrently retained in the active state in VWM is dependent on the interval between them. But the temporal interval has no effect once individuals’ mindset of storage manner has formed.

The different requirements of maintenance could possibly impact the response profile, reflecting the dynamic storage of representations in VWM. Thus, it remains to be explored how the two dynamic arrays are stored in VWM when performing a dynamic recall report task. In addition, given that the fundamental property of the VWM system is limited-capacity, it is still unclear whether the capacity could expand by transferring the currently task-irrelevant stimuli into the passive state. Further studies need to explore these questions to provide new insights of the VWM system.

## Additional information

## Acknowledgement

This research was supported by grants from the National Natural Science Foundation of China (NSFC31970989)

## Author Contributions

Z.L. and J.Z. designed and implemented the experiment; Z.L. and T.L. collected and analyzed the data; Z.L. C.Y. and Q.L. discussed the result of the data and wrote the original draft; Z.L. and C.Y. prepared the figures 1-4; Z.L., C.Y. and Q.L. revised this paper; Q.L. approved the submitted version; all authors reviewed the manuscript.

